# Electrically Contrasting Periodic Polymer Interfaces Guide Neuronal Networks

**DOI:** 10.1101/2025.10.18.683198

**Authors:** Anushka Sarkar, Vanshita Ramsinghani, K. S. Narayan

## Abstract

The nervous system constitutes a highly ordered, integrated network of cells. Understanding this neuroanatomical architecture in vitro is fundamental to elucidating the cellular computations underlying functional network formation. Neuronal connectivity orchestrated through axonal pathfinding, is an interplay of biochemical signals and electromechanical properties of the growth substrate. This study focusses on how neuronal morphology and networking are affected by the periodic, micrometer-scale patterned stripes of two electrically contrasting polymers - poly(vinylidenefluoride-trifluoroethylene) (PVDF-TrFE) and poly(3,4-ethylenedioxythiophene) - poly(styrene sulfonate) (PEDOT:PSS). This periodic confinement provides a length-scale driven cue which unfolds as a self-organized, frequency-modulated spatial phenomenon. Primary cortical cultures on these patterned substrates reveal significant differences with more elaborate network formation on PVDF-TrFE stripe. The features observed at the stripe boundaries, along with the dependence on stripe width, suggest that the neurons exhibit a preference to remain confined within the PVDF-TrFE region, with growth cones deflecting away from the PEDOT:PSS regions. These observations show a promising route for development of a functional template capable of directing axonal growth, synapse formation and neural rewiring in early models of connectivity disorders in vivo.

## 1 Introduction

The human brain comprises approximately 10^11^ neurons interconnected through 10^14^–10^15^ synapses [1], forming a highly complex and hierarchically organized network. Neurons are inherently polarized, comprising the soma for metabolic and transcriptional functions, dendrites for synaptic input integration, and an axon that arises from the axon hillock to transmit signals to distal synaptic targets. During neurodevelopment, axons extend toward their nearest synaptic partners under the guidance of highly dynamic structures at their tips known as growth cones. These sensorimotor organelles characterized by filopodia and lamellipodia [2], actively probe the extracellular milieu and respond to diverse guidance cues – typically chemical (chemotaxis)[3,4], mechanical (durotaxis)[5], and electrical (galvanotaxis)[6–8]. Growth cones transduce these environmental signals through substrate-neuron interactions into intracellular cascades that modulate cytoskeletal dynamics to promote cell adhesion, retraction, polarity, pathfinding and synaptic integration[3–5,9,10]. Beyond axons, the soma itself exhibits mechanosensitive and biochemical responsiveness to the extracellular matrix (ECM), thus anchoring itself to the substrate. [11]

To study such mechanisms in vitro, conventional neural culture platforms (e.g., glass, platinum, silicon) are used – which are rigid and bioinert[12], potentially compromising neuronal adhesion and function. In contrast, emerging polymeric substrates - such as elastomers, hydrogels, thermoplastics, ultrathin dielectrics, and biodegradable fibers - offer enhanced biocompatibility by recapitulating key features of the true neuronal microenvironment. These include mechanical compliance (0.1–4 kPa)[13], biofunctional surface chemistries (e.g., ECM proteins, RGD peptides)[13], micro-topographical cues (2–10 µm grooves)[14], and tunable electrical functionality[15,16]. Neuronal stripe assays and microcontact-printed substrates have long demonstrated that alternating biochemical or topographical cues can impose strong binary preferences on axonal growth. Such substrate patterns, from protein stripes to polymer films, establish that the substrate chemistry and geometry critically shape neurite guidance and network organization via contact guidance [17– 23], which can be mathematically modeled as a drift-diffusion process governed by Langevin or Fokker– Planck equations. This microenvironmental structuring is foundational for engineering biomimetic neural tissues and probing the physics of network self-organization [24,25]. To this, biocompatible conductive and piezoelectric polymers have attracted particular interest due to their ability to modulate neuronal activity through surface charge distribution and electromechanical interactions thus altering neuronal network architecture. Notably, piezoelectric PVDF-TrFE and the mixed ionic–electronic conductor PEDOT:PSS represent two prominent biocompatible platforms in neural interface research [15,26–40], which are the polymers of interest in our study.

PVDF-TrFE exhibits intrinsic electrical activity due to its ferroelectric capacitive “islands” which generate localized electric fields upon mechanical deformation because of its elastic modulus of 4.89 GPa[41]. These fields enhance neuronal attachment, maturation, and synaptic signaling, crucial for neural interface integration and bioelectronic modulation [32,42]. Pristine PEDOT:PSS on the other hand, widely known for its high charge transfer and electrical conductivity, allows for effective electrical signaling and modulation of neural activity. Moderate hydrophilicity (contact angle 50°)[43], high mechanical conformability with a Young’s Modulus between 0.8 and 2.4 GPa[44]; and ability to tailor nano-topography supports long-term neuronal viability and synaptogenesis, especially when coated with adhesion-promoting molecules like poly-D/L-lysine or laminin, fostering functional neuronal networks and reducing glial reactivity [38,45,46].

Independent cell line studies of neuronal connectivity, network formation and morphology have been made on PVDF-TrFE and PEDOT-PSS - making them suitable platforms for advanced neural interfacing and regenerative applications [47,48]. It is interesting to explore the response of neurons to such repetitive patterns as the rationale underlying this approach is based on the principle that neuronal growth and connectivity are influenced by local geometric and material cues, which can be interpreted as frequency modulated spatial features[49]. Such boundary conditions of electro-mechanically contrasting PVDF-TrFE and PEDOT:PSS surfaces of different length scales, have been explored in this study for the first time, to understand the neuroanatomical connectivity and growth dynamics at the junction. Cellular morphometrics address this gap by elucidating the mechanistic relationship between substrate geometry and the generation of specific guidance cues that organize neuronal network architecture.

## 2 Methods

### 2.1 Fabrication Workflow

ITO-coated glass substrates (1.5 × 1.5 cm^2^, Ossila) were sequentially sonicated for 10 minutes each in 2% Extran, deionized water, and IPA:acetone (1:1), followed by cleaning with standard RCA-1 solution (DI H_2_O:NH_4_OH:H_2_O_2_, 5:1:1) and activation in air plasma for 5 minutes. PVDF-TrFE (75:25 molar ratio, PolyK Technologies, USA; 40 mg mL^−1^ in DMSO, SDFCL) was spin-coated onto the activated substrates at 2000 rpm for 30 seconds to yield ∼120 nm films, which were annealed at 120°C for 40 minutes to induce β-phase crystallization. Stripe patterning was achieved using Selective Plasma Assisted Removal (SPAR), wherein PVDF-TrFE films, masked with custom copper patterns, underwent plasma etching to a depth of 120 nm under optimized conditions. PEDOT:PSS (PH 1000, 1.3 wt% aqueous dispersion, Ossila, CAS# 155090-83-8) was filtered using a BD Emerald™ syringe (2 mL) equipped with a PVDF filter (0.45 μm pore size, 13 mm diameter, Axiva) and spin-coated at 1500 rpm for 30 seconds, followed by annealing at 120°C for 30 minutes. Excess PEDOT:PSS was removed from PVDF-TrFE regions, confining the film to the etched stripes. The completed substrates were coated overnight at 37°C with poly-D-lysine (0.1 mg mL^−1^), followed by laminin (5 μg mL^−1^ in PBS, 2 hours at room temperature) immediately before cell seeding.

### 2.2 Characterization of substrates

#### 2.2.1 Atomic Force Microscopy

Non-contact AFM (JPK Instruments, Bruker) images analyzed using JPK Data Processing software revealed the elevated topographical features of PVDF-TrFE with crests up to 50-60 nm high domains and root-mean-square roughness (R_rms_) of 12.93 nm. Meanwhile, the ability of PEDOT:PSS to make smooth granular films at nanoscale was evident by grain size being around 4-5 nm high and R_rms_ of 1.078 nm (Supplementary Figure S1a and S1b). Plasma etching of the polymer enhances the roughness of both polymers by ablating the surfaces. The robust electrostatic interactions between the PEDOT and PSS polymer chains are attributed to lower plasma etching rates of PEDOT:PSS than PVDF-TrFE which is a polycrystalline polymer matrix (Supplementary Figure S1c, S1d and Supplementary Table S1). The interface of the polymers highlights the smoother portion being side filled with PEDOT:PSS and uneven highs comprising polar microdomains of PVDF-TrFE as visualized in AFM scans (Figure 2a and 2b) which corroborate recent findings[50]. Piezo Force Microscopy (PFM) scans of PVDF-TrFE highlight the ferroelectric domain orientation of the surface with respect to applied electric field via the conductive cantilever tip (see Figure 2d and 2e). Kelvin Probe Force Microscopy (KPFM) of PEDOT:PSS provides nanoscale mapping of local surface potential revealing an equipotential polymer profile with a calculated work function of approximately 4.82 eV (see Figure 2g and 2h). Both the studies underline the contrasting electrical neighborhood of the two polymers in context.

**Figure 1.**
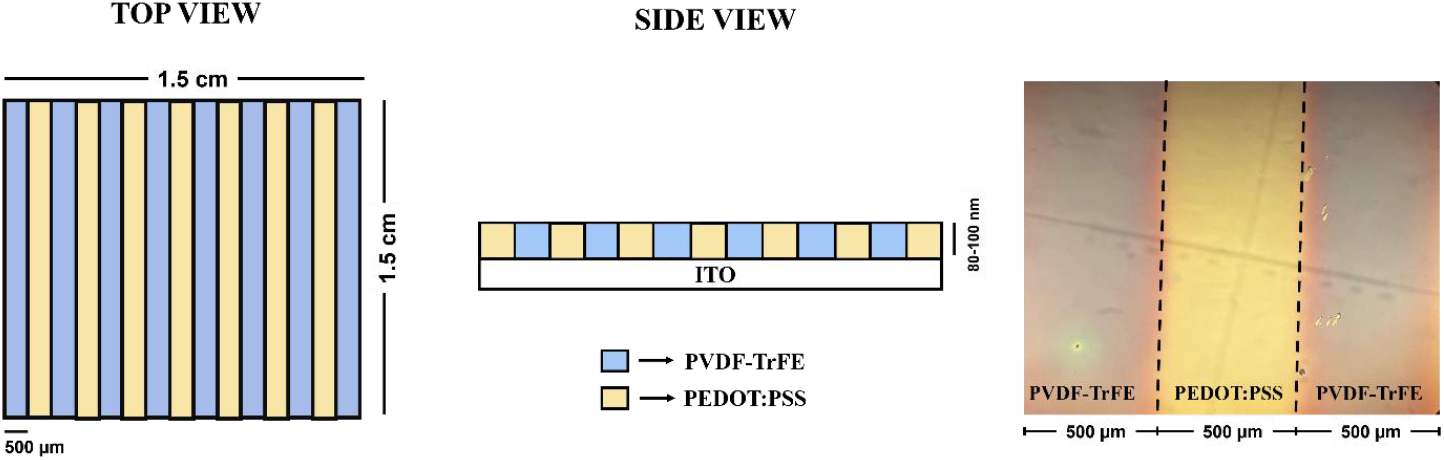
Patterned substrate schematic (left) and stripes seen under brightfield microscope (right) where blue stripe is PVDF-TrFE and yellow stripe is PEDOT:PSS

**Figure 2.**
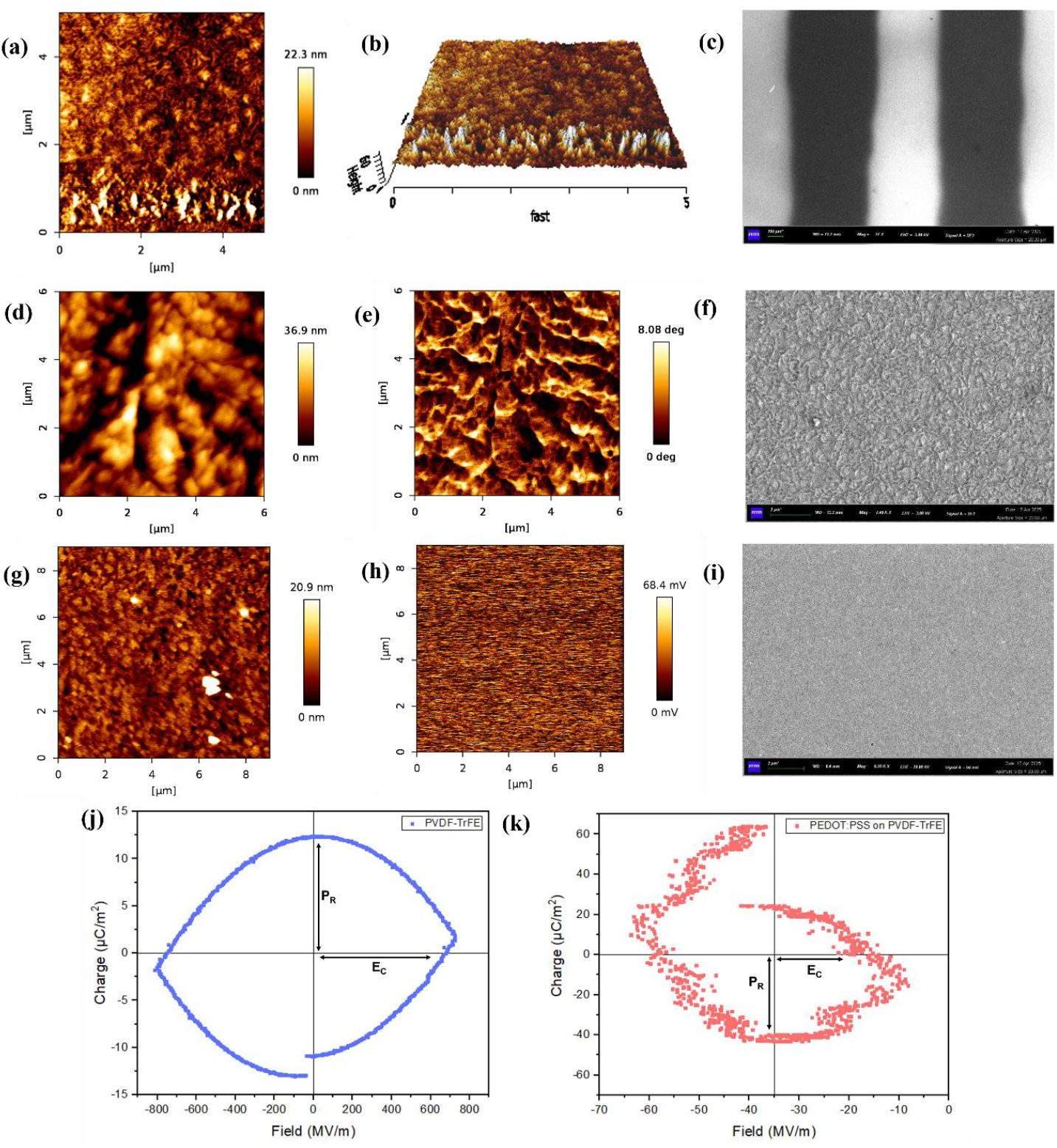
(a) AFM images depicting surface features of the interface region between alternating stripes of PVDF-TrFE and PEDOT:PSS. (b) Three-dimensional reconstruction of the interface highlighting the distinct morphological features. (c) FESEM of juxtaposed microstripes of both polymers where lighter regions corresponding to PEDOT:PSS and dark stripes representing PVDF-TrFE. (d-f) Height trace (R_rms_= 8.21 nm), Phase scan as seen through PFM measurement and FESEM of PVDF-TrFE indicating its ferroelectric domains respectively. (g-i) Height trace (R_rms_= 2.26 nm), Precision 5 scan obtained from KPFM measurement and FESEM of PEDOT:PSS indicating its equipotential granular relief. (j-k) Lateral P-E Loop measurement of PVDF-TrFE (130 nm) and PEDOT:PSS (40 nm) on top of PVDF-TrFE (130 nm) reflecting its in-plane ferroelectric nature arising from the bulk of polymer.

#### 2.2.2 Field-Emission Scanning Electron Microscopy

FESEM of the patterned substrate clearly resolves the distinct lanes of the patterns, light and dark regions being PEDOT:PSS and PVDF-TrFE respectively.(Figure 2c). The scans of PVDF-TrFE show the topographical landscape from a single plane hinting about the ferroelectric domains’ boundaries of the polymer (Figure 2f) that appears as a mosaic of ridges. In its β-phase, PVDF-TrFE is spontaneously polar, and the film self-organizes into 180° or 90° domains separated by charged walls. Spherulitic crystal growth during solvent evaporation further amplifies topography by stacking tilted lamellae, giving the polymer its characteristic undulating relief. Whereas PEDOT:PSS shows a comparatively homogeneous fine-grain landscape due to the presence of phase separated regions of electrically conducting PEDOT and impeding PSS fraction as seen in (Figure 2i).

#### 2.2.3 Polarization-Electric Field

Planar Polarization–Electric field (PE) measurements performed on PVDF-TrFE thin films (130 nm) reveal a predominant in-plane polarization response that is strongly dependent on the applied field direction (Figure 2j). This behavior is indicative of a bulk-driven, field-orientable polarization in the copolymer matrix with Remnant Polarization (P_R_) around 12.3 µC/m^2^ and Coercive field (E_C_) around 680 MV/m. The high dielectric loss and significant leakage currents inherent to PVDF-TrFE contribute to the generation of a non-ideal, lossy shape consistent with high conductivity and incomplete domain switching instead of an ideal rectangular ferroelectric hysteresis. The same measurements were performed on the PVDF-TrFE substrate with PEDOT:PSS coated on top (40 nm) which changes the boundary conditions from previous system due to addition of an iso-electric thin layer with finite resistance on top of dielectric bulk of PVDF-TrFE (Figure 2k). The nature of the curve is analogous to ideal PVDF-TrFE but quite scattered and dependent on the applied field but with a shifted bias voltage. Remnant polarization is significantly higher about -41.2 µC/m^2^ with a small coercive field around -20 MV/m.

#### 2.2.4 Contact Angle

Contact angle measurements were conducted to evaluate the wettability of polymer substrates, with a 10 µL water droplet placed on each surface and ITO used as a reference. PVDF-TrFE exhibited higher contact angles than PEDOT:PSS, reflecting its hydrophobicity (Supplementary Figure S2). The lower contact angle of PEDOT:PSS is attributed to the hydrophilic PSS^-^ counterion, which promotes uniform water spreading. Plasma etching increased the hydrophilicity of both substrates by enhancing surface energy and roughness. This effect was more pronounced for PVDF-TrFE, which showed a substantial decrease in contact angle, likely due to its greater susceptibility to oxidative surface modification, while PEDOT:PSS showed only modest changes, consistent with its already hydrophilic character and its higher etching rate. Values are indicated in Supplementary Table S2.

#### 2.2.5 Electrical Characterization

##### PEDOT:PSS

The PEDOT:PSS surface is conductive, with electrical transport occurring through the PEDOT domains, which effectively navigate around the insulating PSS regions. It is to be noted the PSS barriers hamper the vertical electrical transport across the film thickness significantly as compared to the surface transport. The I–V behavior of PEDOT:PSS in the surface configuration is linear with conductivity in range of ∼ 6.6 S/cm. It is known that this conductivity can be further controlled by secondary dopants and film formation processes (Supplementary Figure S3a)[51]. In the present study, the PEDOT:PSS film was annealed prior to surface treatment to optimize its conductive properties and surface characteristics before neuron seeding.

##### PVDF-TrFE

PVDF-TrFE films exhibit insulating behavior with a DC σ < 10^-5^ S/cm and their nonlinear current-voltage characteristics support this observation (Supplementary Figure S3b). AC measurements across varying frequencies confirm the characteristic capacitive response of the films, with a high dielectric constant (∼8) observed at low frequencies due to efficient reorientation of molecular dipoles and domains in response to slowly changing electric fields The dielectric constant decreases as frequency increases, reflecting dipole dispersion within the polymer matrix. ε(ω) studies of PVDF-TrFE have been extensively studied and are well understood in terms of multiple relaxation models. At high ω, the polarization mechanisms are unable to keep pace with the rapidly oscillating field, resulting in a decrease in the dielectric constant (Supplementary Figure S3c). Correspondingly, the impedance of material increases at higher frequencies due to the reduced contribution of dipolar polarization. [52]

### 2.3 Primary Neuron Culture

Primary neuron cultures are ideal biointerfaces next to in-vivo experiments, because they maintain native ion channels and synaptic plasticity, closely mimicking in vivo brain physiology. This makes them the gold standard for initial testing of brain-machine interfaces and device compatibility. The complex neurite outgrowth and network formation achieved provide realistic functional readouts, improving biointerface assessment accuracy. Therefore, primary cortical cultures were chosen for their physiological relevance and robust performance.

Primary cortical neurons were isolated from P0–P1 C57BL/6 or BALB/c mice and cultured in a defined, serum-free medium. Brains were dissected in ice-cold Hibernate™-A medium (Cat. No. A1247501, Gibco™) supplemented with 1% GlutaMAX™ (Cat. No. 35050061), 1% Penicillin–Streptomycin (Cat. No. 15140122), and 2% B-27™ Supplement (Cat. No. 17504044), filtered through a 0.2 µm PTFE membrane (Axiva Filters). Cortices were isolated, cleared of meninges and white matter, minced (∼1 mm^3^), and enzymatically dissociated using 0.25% Papain (Cat. No. P4762, Sigma-Aldrich, Merck) and DNase I (Cat. No. M0303L, New England Biolabs) at 37 °C for 15 min. The suspension was triturated, filtered (0.2 mm mesh), centrifuged (5 min, 80 × g), and resuspended in Neurobasal™-A medium (Cat. No. 10888022) supplemented as above. Cells (∼2 × 10^5^ cells/35 mm dish) were plated on Poly-D-Lysine–coated (Cat. No. A3890401) substrates and maintained at 37 °C, 5% CO_2_ with half-media changes every 3 days. On DIV3, Cytarabine (Ara-C, 10 mM) was added to suppress glial proliferation, if any. Detailed protocol as a schematic in Supplementary Figure S4.

### 2.4 Data Analysis

Neuronal outgrowth was monitored under phase-contrast microscopy from DIV 0 to DIV 14 using Capture 2.4 software and analyzed in Fiji ImageJ using Simple Neurite Tracer (SNT) plugin. Manual or semi-automated tracing was performed to accurately delineate neurite paths, while thresholding and skeletonization provided objective identification and measurement of neurite architecture. All analyses were performed using five independent biological replicates, each comprising two technical replicates. For each technical replicate, data were collected from 10–12 individual stripes per substrate. This multi-tiered experimental design ensured robust statistical power for quantitative comparison of neuronal morphologies and network dynamics across the different substrate conditions.

## 3 Results

Our study reveals that both PVDF-TrFE and PEDOT:PSS independently support neuronal adhesion and neurite outgrowth as illustrated in Figure 3a & 3b - consistent with earlier reports highlighting the biocompatibility and electroactive nature of the substrates [15,38]. Interestingly, neurons cultured on glass -commonly used as a control in in-vitro studies - exhibited poor adhesion, sparse neurite extension, and limited branching (Figure 3c). Such behavior aligns with established mechanotransductive responses of neurons, where stiff substrates such as glass or highly cross-linked polymers activate RhoA/ROCK signaling, reducing forward neurite extension and promoting local branching[53]. It was also observed that neuronal networks were more elaborate on PVDF-TrFE control compared to PEDOT:PSS control (Figure 3d), as the network matures slower on PEDOT:PSS. Collectively, these findings highlight the intrinsic potential of electroactive polymers like PVDF-TrFE and PEDOT:PSS as favorable substrates for neuronal culture, supporting cell adhesion and neurite outgrowth independently of any imposed patterning or topographical modifications.

**Figure 3.**
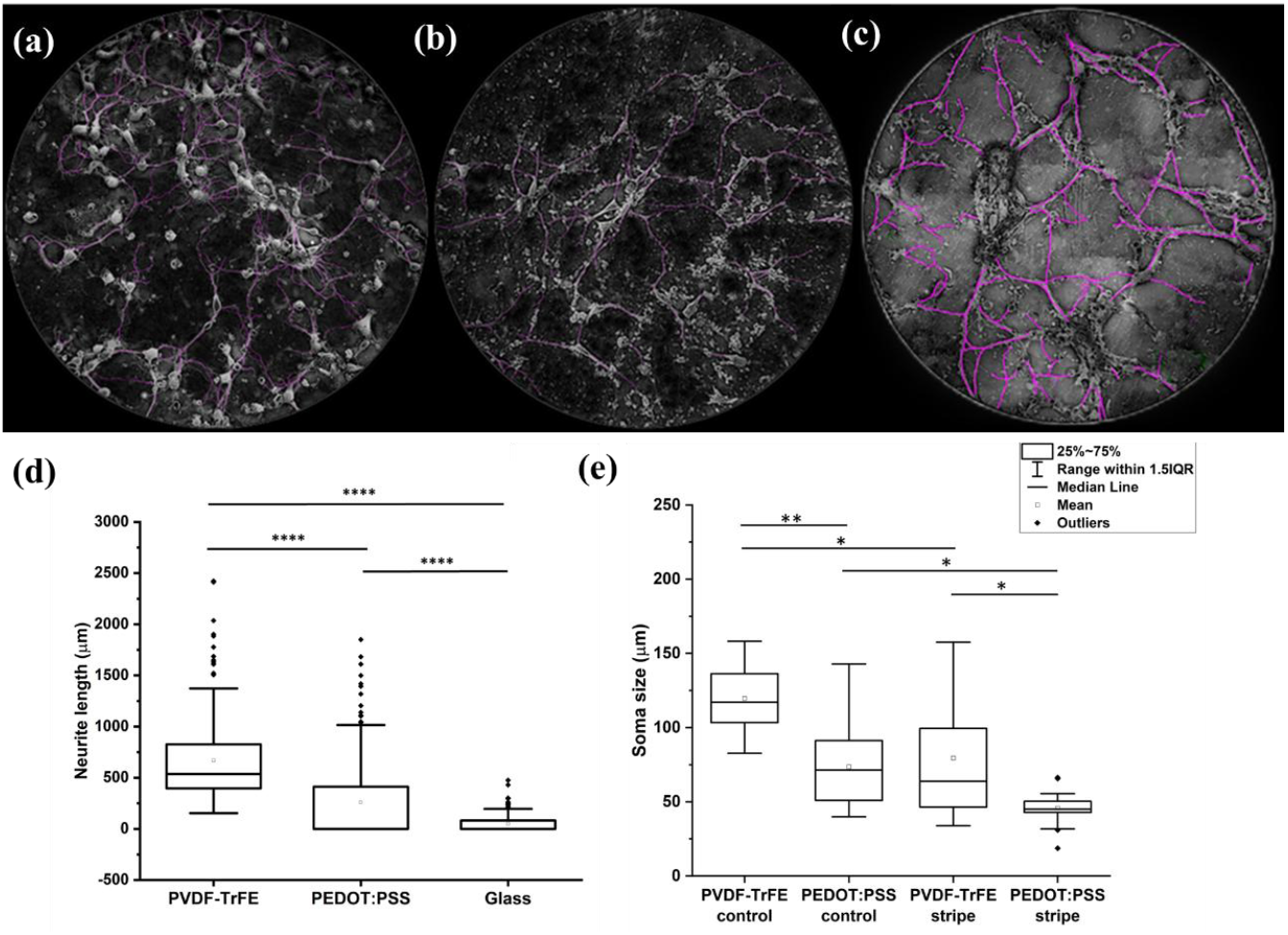
Neurite length distribution on different substrates. Representative phase-contrast images (ImageJ neurite-trace overlays) of culture on (a) PVDF-TrFE, (b) PEDOT:PSS and (c) PDL–glass substrates at DIV 10. (d) Quantitative analysis of neurite length on each substrate, presented as box-and-whisker plots; statistical significance is indicated (***p < 0.0001). Both polymeric substrates supported substantially longer neurite extension compared to glass. Among the polymers, PVDF-TrFE promoted the greatest neurite outgrowth. (e) Comparing soma sizes of neurons cultured on PVDF-TrFE and PEDOT:PSS substrates under control and stripe-patterned conditions. Soma size is significantly reduced on PEDOT:PSS stripes and control compared to all other groups (*p < 0.05, **p < 0.01). Scale: 3.165 px/µm.

Further morphometric analyses of neuronal cultures grown on striped substrates revealed substantially increased neurite branching complexity and enhanced alignment on PVDF-TrFE regions relative to PEDOT:PSS. Quantitative assessments on the interface demonstrated a statistically significant preference for neurite localization on PVDF-TrFE stripes, with an average of 65.2% of neurites confined to PVDF-TrFE regions versus 34.8% extending into PEDOT:PSS domains (Figure 4f), directly confirming robust, material-specific guidance. Additionally, neuronal cultures on PVDF-TrFE exhibited increased soma size (Figure 4e), greater neuron density, and elongated neurites relative to those on PEDOT:PSS, as shown in Figure 4c and corroborated by control experiments. Collectively, these results highlight the fundamental influence of substrate material properties in directing neuronal morphology, spatial organization, and maturation trajectories.

**Figure 4.**
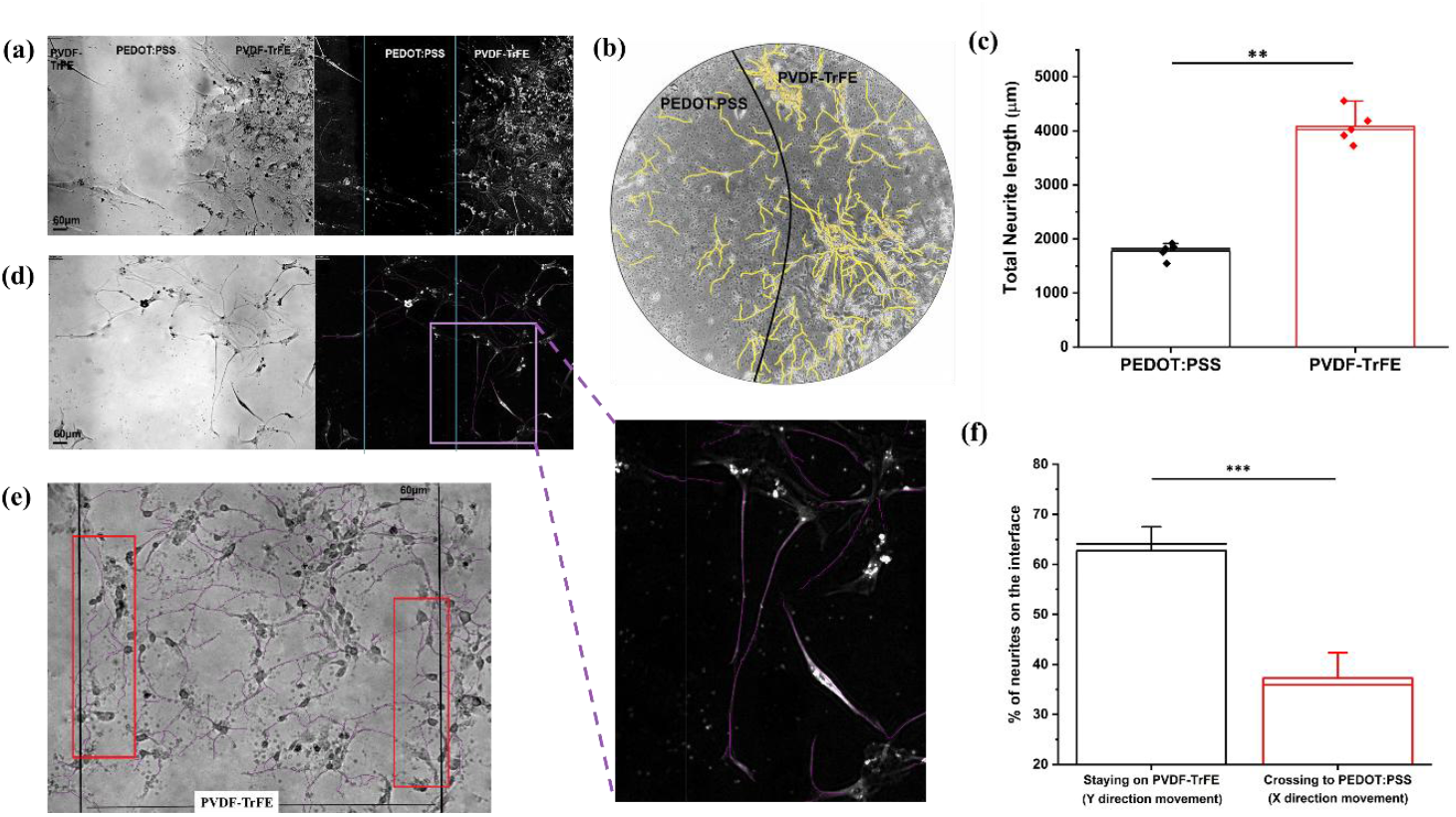
(a) Representative brightfield and neurite-traced images of neuron culture on linear stripe pattern of 500 µm showing higher neuron density and neurite length on PVDF-TrFE. (b) Concentric circular patterns of diameter = 1000 µm composed of alternating regions of PVDF-TrFE and PEDOT:PSS in neurite traced image (Black curved lines indicate the demarcation between the polymers). Neuronal cultures exhibit higher cell density and pronounced neurite curvature within the PVDF-TrFE domains, consistent with the directional growth observed on vertically aligned stripe configurations. (c) Quantification of total neurite length reveals a significant increase within PVDF-TrFE domains compared to PEDOT:PSS. (d-e) The guidance of neurite as observed (inset) on the polymer interface on stripe pattern of 500 µm and 1000 µm respectively. (f) Statistical analysis of neurites curving at the interface by plotting % of neurites crossing boundary versus staying on PVDF-TrFE polymer. (f) Analysis of neurite trajectory at the interface demonstrates a markedly higher proportion of neurites remain within PVDF-TrFE (Y direction) relative to those crossing into PEDOT:PSS (X direction) - suggestive of material-guided navigation (**P < 0.01, ***P < 0.001; one-way ANOVA, post-hoc: Tukey’s HSD). Scale bar: 60 µm. Errors bars: SD.

## 4 Discussion

The substrate provides a periodically distinct surface field to the neurons arising from the intrinsic electrical nature of the polymers. The data consistently indicate that the ferroelectric PVDF–TrFE surface serves as a localized charged anchor, stabilizing neuronal somata and guiding neurite outgrowth due to its slightly higher stiffness than PEDOT:PSS [28,54]. The PVDF-TrFE domains provide a distinct combination of topographical and ferroelectric surface properties, which are hypothesized to facilitate neurite extension by modulating cell adhesion, membrane tension, and intracellular signaling pathways. Neurons cultured on PVDF–TrFE receive dynamic electromechanical cues that enhance neurite extension, modulate calcium oscillations, and bias axonal alignment. The microscopic electric cues to cells have been attributed to the intrinsic polar domains [55]. The contribution of the electric field at the surface arises from the depolarization field (see Equation 1) and surface bound charges (see Equation 2) and the polarization discontinuities at the air-polymer interface. [56]

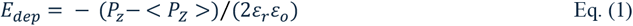

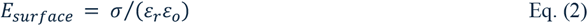

Where, Pz and <Pz> is the local and averaged ferroelectric polarization over the film area, *ε*_0_ and *ε*_r_ are the vacuum permittivity and static dielectric constant of the material and surface charge density is 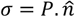 due to polarization along normal vector, respectively. In contrast, the electrically conductive PEDOT:PSS layer acts as a passive, low-impedance substrate that supports neuronal growth. The electrical landscape is influenced by the electrical double layer (EDL) formed in the micellar structure between phase separated PEDOT and PSS^-^ due to their charge difference. The distributed clusters of PEDOT core covered by PSS^-^ shell is the source of volumetric capacitance which along with the work function (Φ) is contributing to the surface fields.[57]

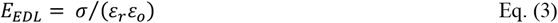

Here, surface charge density *σ* = *e*(*c*^*+*^ *-c*^*-*^) is due to hole concentration (*c*^*+*^) in PEDOT and concentration of sulfonate ions (*c*^*-*^) in the PSS chain which contributes to electric field (see equation 3). The PEDOT:PSS film surface offers heterogenous variations of the surface conductance at nanoscale. These localized fields can influence neuronal navigation by acting as an electrostatic arena differentially steering growth cone.

Further, the growth cones on axons extend like hooks ready to anchor themselves to the substrate or nearest synaptic partners. The surface topography of PVDF-TrFE assists in docking of soma on initial days of seeding while they also migrate when tension develops between axons of nearby neurons pulling them together. Hence, resulting in higher

density on textured PVDF-TrFE surface than smoother surface of PEDOT:PSS [55] Such guidance may manifest as neurite deflection or curvature away from regions of like charges, consistent with electrostatic repulsion phenomena (See Figure 4d - inset). Careful analysis of substrate reveals that there is a thin residual layer of PEDOT:PSS on PVDF-TrFE layer as a remnant of the plasma etching process. However this does not affect the underlying elastic and ferroelectric properties of PVDF-TrFE, as confirmed by P-E loop measurements (Figure 2j, k). The bending of neurites is an inherent signature which is consistently observed at the ferroelectric stripe-conducting stripe boundaries. A significant neuron growth was also observed along the perimeter of the substrate. It is to be noted that the perimeter region is slightly thicker and of heterogeneous composition that promotes neuron growth (Supplementary Figure S5).

PVDF-TrFE has intrinsic piezoelectricity which means as the neurons exert traction forces on the substrate, localized voltage pulses are generated that open voltage-gated Ca^2+^ channels in the growth cone membrane [54]; these Ca^2+^ microdomains drive actin polymerization and filopodial stabilization more robustly than on nonpiezoelectric PEDOT:PSS - which might aid to the guiding of neurons[58]. Also, the relatively high elastic modulus (∼4.89 GPa) of PVDF-TrFE [41] places growth cones into the “load and fail” regime of the motor–clutch system, producing greater traction forces and longer adhesion lifetimes that could favor neurite initiation and cell spreading, whereas the softer PEDOT:PSS substrate elicits faster retrograde flow with lower traction[15]. From the wettability point of view, the more hydrophobic PVDF-TrFE surface adsorbs ECM proteins (e.g., fibronectin) in conformations that enhance integrin α5β1 clustering and downstream FAK–RhoA signaling, promoting stable focal adhesions and increased neuronal attachment [59], while the hydrophilic PEDOT:PSS yields less favorable protein conformations and reduced integrin engagement. Thus, the ability of PVDF-TrFE to retain adsorbed neurotrophins (BDNF, NGF) via electrostatic interactions may amplify local Trk-receptor signaling, further boost directed outgrowth and branching density of the growth cones compared to PEDOT:PSS.

The recurring boundary conditions are explicitly dependent on the stripe dimensions. An optimal microstripe width can significantly promote neurite alignment and network connectivity. Hence, neuronal cultures were performed as a function of different stripe widths of 300 µm, 500 µm, 700 µm and 1000 µm and it is observed that a length scale ≈ 500 µm is effective in curving the neurites (Figure 4a). This might be because of three factors – it balances confinement to enhance degrees of freedom for natural neurite branching and alignment, supports stable gradients of guidance molecules, and enables proper cytoskeletal organization. This size facilitates cell adhesion and mimics natural tissue scales, promoting effective directional neurite growth and network formation. Around 300 µm, there was highest branching density with dense, highly branched neurites but limited neurite extension due to spatial constraints. The reason for this could be explained by homogeneity of surface fields due to small confinement length. Whereas in larger widths of 700 - 1000 µm, neurites presented moderate degrees of branching; but their orientation displayed reduced spatial alignment, with axonal trajectories becoming irregular and exhibiting inconsistent growth patterns. The neuronal network formation on individual PEDOT:PSS/PVDF-TrFE stripe of this length scale is similar to the respective polymer control except for the guiding phenomenon seen at the junction which is not visible on controls due to absence of periodic polymer interface.

To introduce other parameters which might play a role but were difficult to account for experimentally; a dual-model computational framework was employed grounded in established theories of neuronal morphogenesis [60,61]. The primary model was a detailed, agent-based simulation in which neurite extension was represented as a biased random walk. The direction of neurite growth was subject to a tunable angular bias that diminished with increasing stripe width, reflecting the reduced efficiency of topographical guidance on wider substrates. Significant angular deviations were classified as branching events, in accordance with the stochastic branching paradigm employed in previous kinetic and biophysical models. A secondary, simplified stochastic model was developed to independently validate the empirically observed trade-off between directional alignment and branching frequency across stripe widths ranging from 300 to 1000 µm. For each parameter set, we computed:

- **Mean Directional Alignment:** Quantified as the average of cos^2^θ, where θ is the angle between the neurite trajectory and the axis of the microstripe, in line with conventions in the computational modelling of neurite guidance.
- **Branching Density:** Defined as the average number of large-angular-deflection events per simulated neurite, parameterizing network complexity.

Both metrics were normalized to the unit interval to facilitate comparative analysis. The optimal stripe width was operationally defined as the intersection point of the normalized alignment and branching density curves, corresponding to the regime in which substrate cues optimize alignment without excessively constraining network complexity. Our simulations identified this intersection consistently within the 500-630 µm range (∼42 runs) (Figure 5b). This result is consonant with theoretical predictions based on coupled differential equations for neurite outgrowth and stochastic models of branching, which also concretes our experimental observations, which indicate that optimal substrate feature size aligns with the intrinsic exploratory length scale of neurites. While a closed-form analytical solution for the optimal width is elusive due to the stochastic and nonlinear nature of neurite dynamics, our parameter sweep, and intersection analysis provide a robust computational method for substrate design optimization.

**Figure 5.**
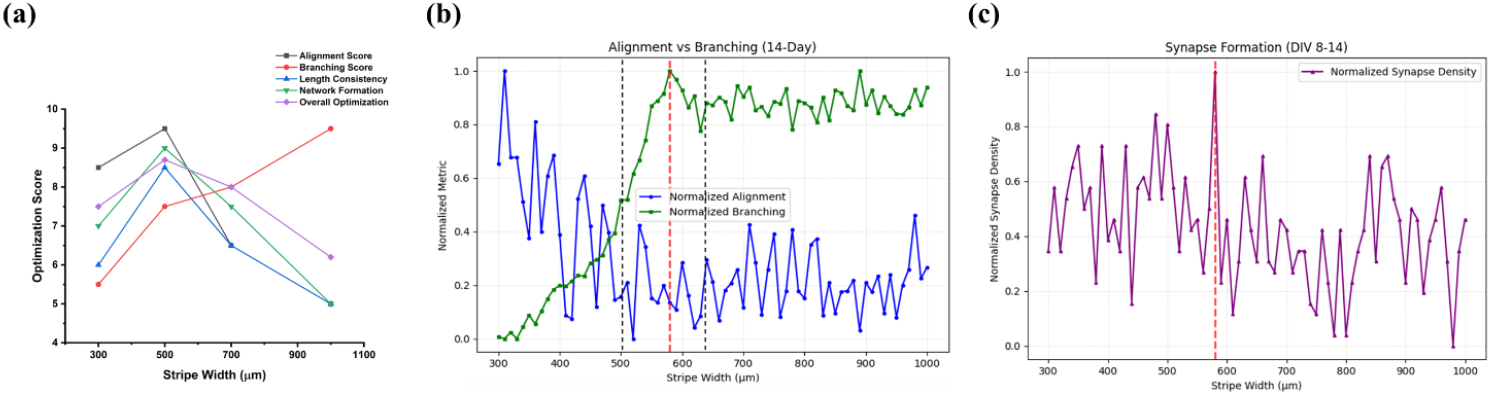
(a) Experimentally derived optimization scores for different network parameters (alignment, branching, length consistency, network formation, and overall optimization) as a function of stripe width – with a peak at 500 µm. (b) Simulation-derived optimal stripe width for neurite alignment, branching, and synapse formation across varying microstripe widths. The intersection of alignment and branching curves occurs at approximately 580 µm (with an optimum range from 500 - 630µm), correlating with composite optimization score. Guidance effectiveness varies over growth phases, with active outgrowth showing highest influence. (c) Normalized synapse density as a function of stripe width also reveals a peak in synaptogenesis around 600 μm, indicating optimal microenvironment dimensions for synapse formation during DIV 8–14.

Introducing spatial modulation in surface fields by juxtaposing electro-mechanically contrasting polymer creates localized fields at the junction that can influence neuronal navigation by acting as an electrostatic barrier differentially steering the growth cone. The magnitude of the electric field discontinuity at the polymer junction, combined with the physical stripe width, defines the effective barrier height and width analogous to a periodic potential well. In this framework, when the energy required for a neurite to overcome the junctional barrier exceeds the energy needed to maintain alignment along the stripe, growth cones favor retention within the same polymer domain, resulting in neurites orienting parallel to the stripe axis (Y direction). PVDF-TrFE emerges as an ideal substrate for supporting complex and elaborate neuronal network formation, both in unpatterned (control) and stripe-patterned configurations. In contrast, PEDOT:PSS displays significant differences in neuronal branching, neurite length, and size between its control and patterned states, making it a practical tunable parameter for modulating the barrier width.

In control experiments using uniform films of PEDOT:PSS and PVDF-TrFE, neuronal outgrowth exhibits a two-dimensional spatial distribution i.e. isotropic both in X and Y direction (Figure 3a, 3b). However, introducing periodicity in the substrate architecture elicits an emergent, stripe-width-dependent organization that transforms the growth problem into a one-dimensional anisotropic system (X being the anisotropy direction and Y being direction along polymer axis statistically favored) constrained by periodic boundaries. This emphasizes a key feature of our system that - accessing a matrix of repetitive interfaces is not equivalent to inspecting a single junction below characteristic length-scale. Neurite guidance is common at all interfaces, but stripe widths exceeding critical thresholds act as independent junctions. In contrast, length-scale below this threshold neurite guidance involves correlated interactions across multiple adjacent interfaces, producing a collective network response directly tied to the spatial frequency of the substrate. This capacity of neurons to detect periodic spatial modulation indicates that their complex patterns of growth encode hidden frequency components intrinsic to their collective architectures. This can be resolved and interpreted through Fourier-based analytical methodologies to interpret emergent anisotropy and the fundamental relationship between neuronal organization and substrate topography [49].

## 5 Conclusion

Previous studies have demonstrated the use of PVDF-based scaffolds for neuronal regeneration and in other cell types[32,36,37,42,62,63]. But this electroactive interface of PEDOT:PSS and PVDF-TrFE offers the potential to guide neurite outgrowth, thereby enabling the strategic routing of neuronal processes. The findings establish a proof-of-concept bio-interfacing model in which spatially patterned stripes of PVDF-TrFE and PEDOT:PSS exert differential guidance cues for neuronal growth. Specifically, the ferroelectric PVDF-TrFE lanes serve as preferential conduits for axonal elongation, likely by favorable electrochemical interactions at the growth cones which promote more stable anchoring of axons, thereby enabling efficient navigation toward neighboring targets, whereas PEDOT:PSS - despite being softer and electroconductive - tends to support less robust neurite outgrowth, likely due to weaker anchoring and less effective guidance cues. [64–66]. The interfacial boundary regions act as barriers or decision points, where axons frequently either reorient their trajectory or initiate collateral branching on the PVDF-TrFE side. By combining the ferroelectricity of PVDF-TrFE with the mixed ionic–electronic conductivity of PEDOT:PSS in a spatially patterned manner, we demonstrate a material-based strategy capable of biasing axonal decision-making in a predictable fashion. Such interfaces may be particularly applicable in early developmental models of neuronal connectivity disorders or mild injury, over denervated cortical or spinal regions where the capacity for synaptogenesis and network remodeling remains high. Although direct implantation of this guidance process remains to be explored, this exercise provides fundamental evidence of a scalable system for controlled network rewiring and synaptic integration.

## Ethics Statement

All studies were performed in accordance with the guidelines of the Institutional Animal Ethics Committee (IAEC) and the Institutional Bioethics and Bio-Safety Committee, Jawaharlal Nehru Centre for Advanced Scientific Research. All protocols and experiments were approved by the IAEC (Proposal No. KSN 003).

## Data Availability

Data supporting the findings of this research will be made available upon reasonable request.

## Code availability

Simulations were performed using custom code written in Python that is available upon reasonable request.

## Author Contributions (CRediT)

**Anushka Sarkar:** Data curation, Formal analysis, Investigation, Methodology, Validation, Visualization, Writing – original draft, Writing – review & editing. **Vanshita Ramsinghani:** Investigation, Methodology, Validation, Writing – review & editing. **K. S. Narayan:** Conceptualization, Funding acquisition, Project administration, Resources, Validation, Software, Supervision.

## Acknowledgements

The authors would like to acknowledge Ravi Manjithaya of Molecular Biology and Genetics Unit, JNCASR for Cell Culture Facility and RG Prakash for Animal House facilities. We also thank the Chemistry and Physics of Materials Unit, JNCASR, for access to central facilities, Durgesh Gaikwad of IISER Bhopal, for assistance with data analysis and Ashar A Z for PFM imaging of PVDF-TrFE films.

