## Supplementary File for "Electrically Contrasting Periodic Polymer Interfaces Guide Neuronal Networks"

**Supplementary Figures**

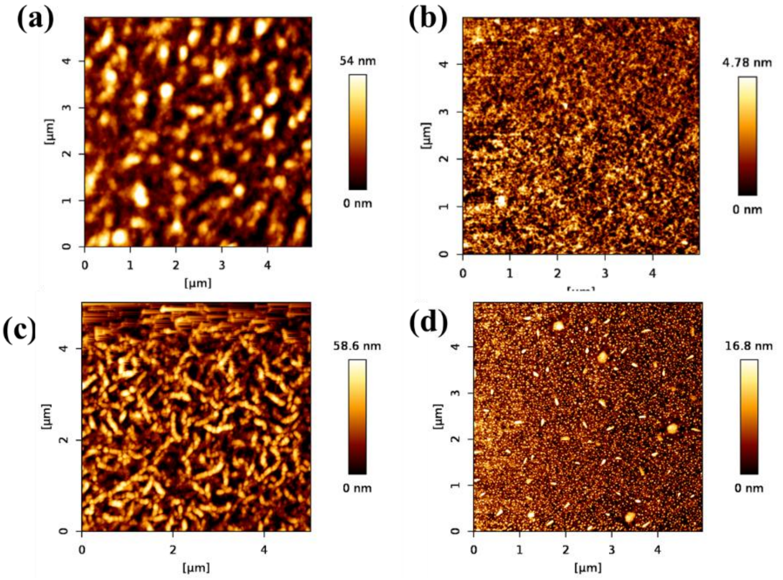


**Figure S1**. Atomic Force Microscopy (AFM) images depicting polymer morphology of (a) PVDF-TrFE ($R_{rms}$= 12.93 nm) and (b) PEDOT:PSS ($R_{rms}$= 1.073 nm). Plasma etching of 6 mins increases the surface roughness of (c) PVDF-TrFE ($R_{rms}$= 13.74 nm), (b) PEDOT:PSS ($R_{rms}$= 3.8 nm)

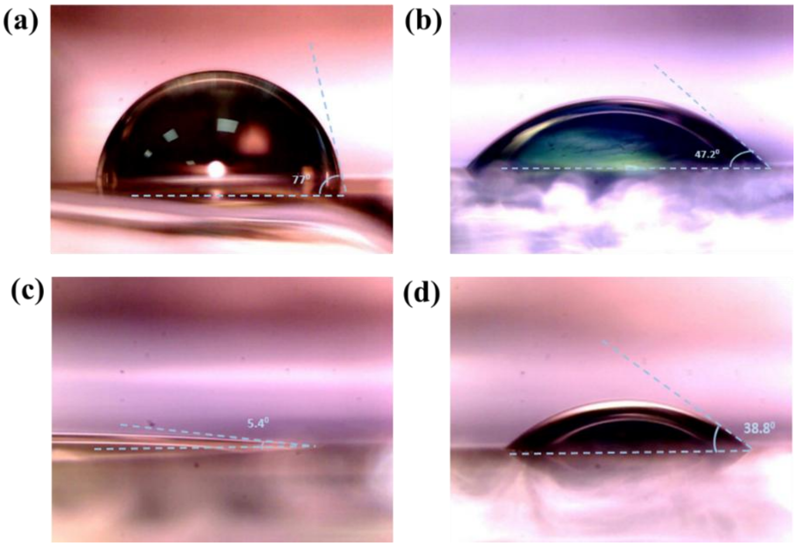


**Figure S2.** Contact angle measurements describing the wettability of (a) PVDF-TrFE and (b) PEDOT:PSS before etching; (c) PVDF-TrFE and (d) PEDOT-PSS post etching

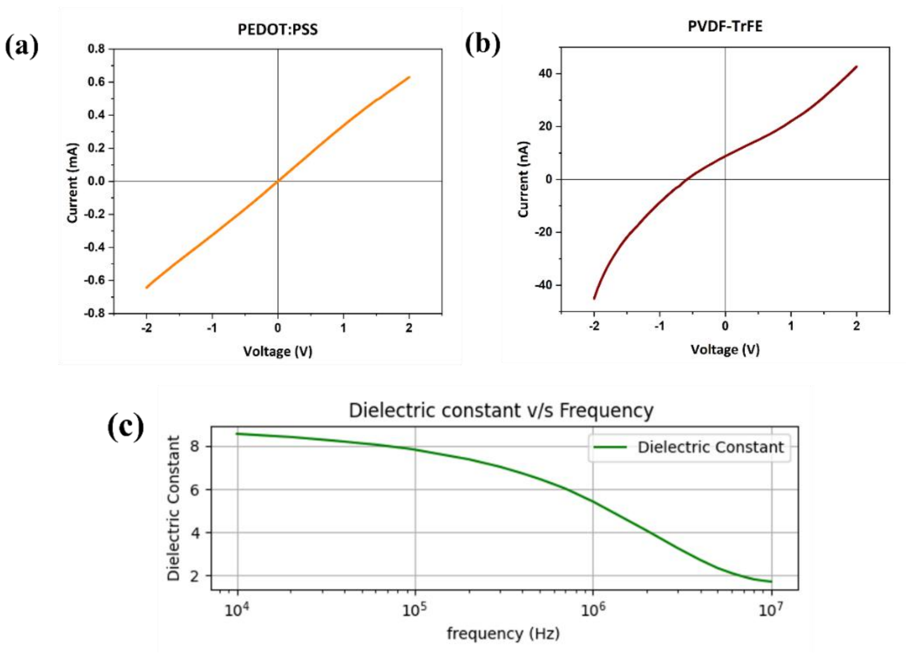


**Figure S3.** (a) I-V measurement of PEDOT:PSS indicating its linearly conducting behaviour. (b) I-V measurement of PVDF-TrFE showing its insulating behaviour with current values in nA. (c) The plot of dielectric constant versus frequency for PVDF-TrFE exhibits characteristics typical of capacitive behaviour.

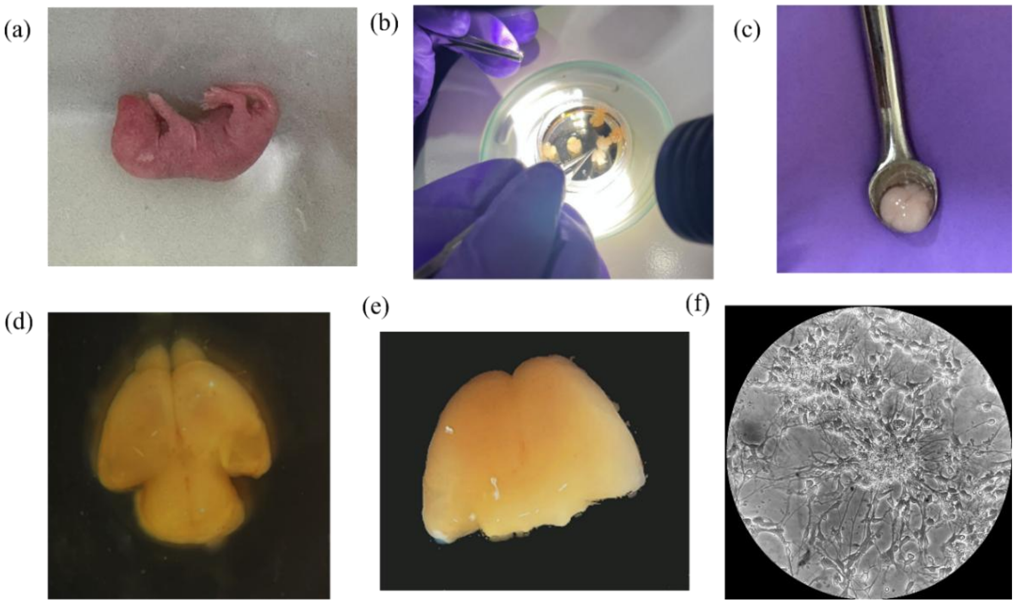


**Figure S4.** Mouse Brain Dissection and Neuron Extraction: Top row (left to right): (a) Postnatal day 1 (P1) mouse pup used for brain dissection; (b) dissection of the brain under a stereomicroscope in ice-cold Hibernate-A medium; (c) isolated intact brain tissue placed in a sterile spoon for transfer. Bottom row (left to right): (d) dorsal view of the freshly dissected brain showing both hemispheres and olfactory bulbs; (e) dorsal view of the same brain's cortex; (f) representative phase contrast image of the neuronal culture on DIV 7.

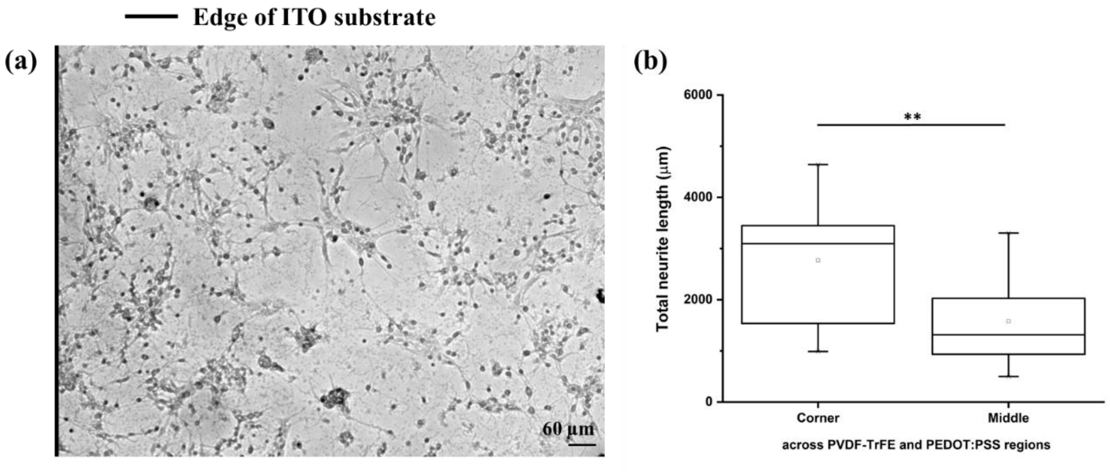


**Figure S5.** (a) Representative phase contrast micrograph showing enhanced neuronal network morphology at the edge of the ITO substrate. Scale bar: 60 µm. (b) Box plot illustrating total neurite length measured in corner and middle regions across PVDF-TrFE and PEDOT:PSS substrates. Neurite extension is significantly greater at corners (where overlap of polymers is present) compared to middle regions (**$p < 0.01$).

**Supplementary Tables**

| **Polymer** | **RMS Roughness (nm)** | **Height (nm)** |
|---|---|---|
| **PEDOT-PSS** | 1.073 | 50.41 ± 10 |
| **PEDOT-PSS Post Etching 6 mins** | 3.8 | 30.51 ± 6 |
| **PVDF-TrFE** | 12.93 | 100.15 ± 15 |
| **PVDF-TrFE Post Etching 6 mins** | 13.74 | 47.73 ± 8 |
| **PEDOT-PSS on Alternating Stripes** | 4.69 | 70.19 ± 4 |
| **PEDOT-PSS Spin coated on PVDF-TrFE Stripes** | 11.47 | 140.66 ± 11 |

**Table S1.** Estimated Roughness and Thickness measurement of Polymers measured by AFM between steps in the fabrication process

| **Polymer** | **Contact Angle (deg)** |
|---|---|
| **ITO** | 37.57 ± 0.94 |
| **PVDF-TrFE** | 77.78 ± 1.16 |
| **PVDF-TrFE Etched 5 mins** | 6.85 ± 1.09 |
| **PEDOT:PSS** | 55.54 ± 5.14 |
| **PEDOT:PSS Etched 5 mins** | 39.17 ± 1.33 |

**Table S2.** Contact angle of PVDF-TrFE and PEDOT-PSS pre and post etching
